# From neuropeptide and receptor annotation to ligand-receptor pairing: a sequence- and structure-based framework for mapping the neuropeptide-receptor interactome in *Gryllus bimaculatus*

**DOI:** 10.64898/2026.08.28.747697

**Authors:** Febrina Margaretha, Mika Sakamoto, Hitomi Seike, Shinji Nagata, Yasukazu Nakamura, Takako Mochizuki

## Abstract

Neuropeptides and their G protein-coupled receptors (GPCRs) control much of insect physiology and behaviour, but in *Gryllus bimaculatus*, an emerging model and edible insect, receptor sequence similarity hinders the mapping of which peptide each GPCR activates. We re-annotated a chromosome-scale genome (BUSCO 95.3%, from 86.7% on insecta_odb12) with comprehensive curation of 48 neuropeptide precursor families (51 loci, including seven not previously identified) and 134 candidate GPCRs (66 rhodopsin-class, 68 secretin-class), providing a near complete neuropeptide-receptor interactome catalogue. We modelled all 15,946 peptide-receptor pairs with AlphaFold3 and Boltz-2 and scored each interface with pLDDT and ipSAE. Ranking these scores, and cross-checking the top candidate for each family against a receptor phylogeny of known ligand specificity, gave a confident, phylogenetically related receptor for 27 of 35 curated receptor groups. These matches confirm the structural scorings with existing deorphanization data and propose receptors for peptides with no prior functional evidence. The annotation, curated peptide and receptor sets, and ranked complexes are available through CricketBase (https://cricket.annotation.jp), a genome browser with a structure viewer of peptide-receptor complexes, providing a resource for *G. bimaculatus* endocrinology and a workflow to deorphanize GPCRs in other non-model insects.

**Graphical Abstract:** Figure 2.

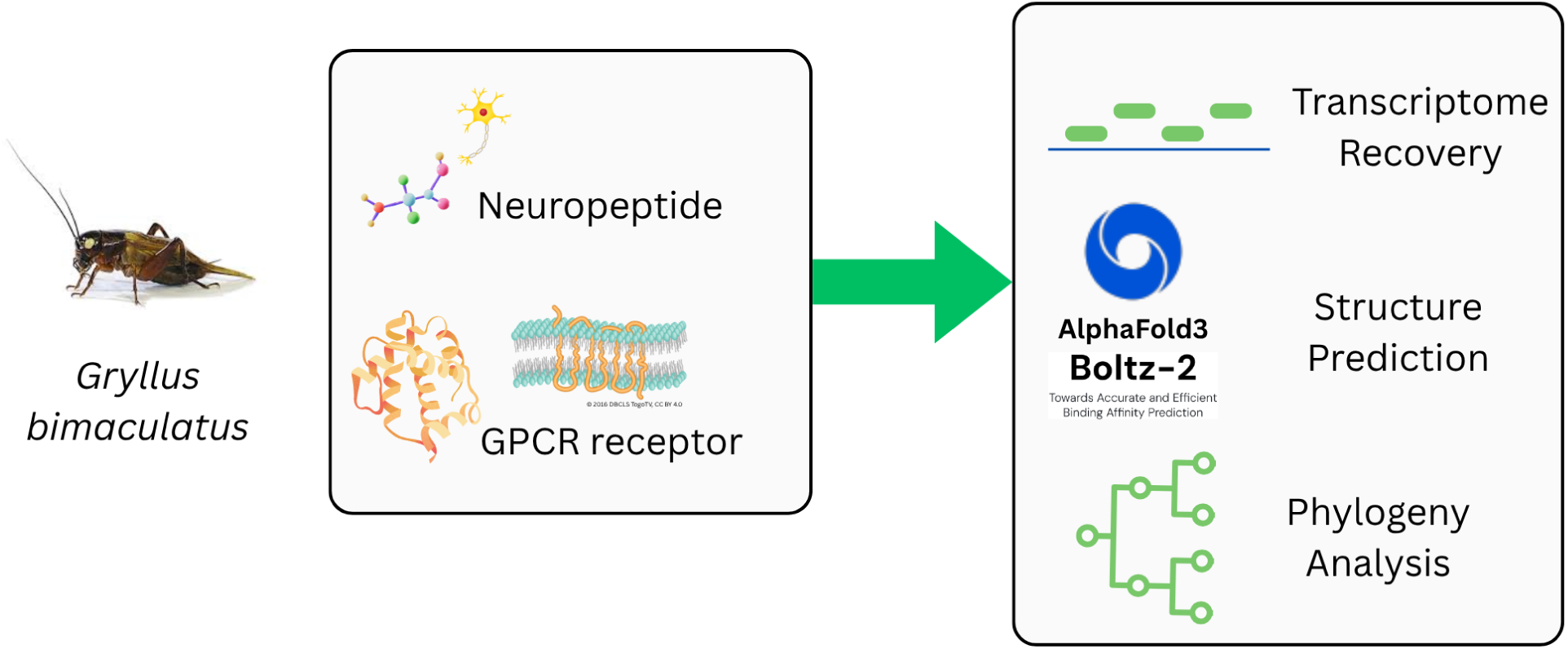

## Introduction

Advances in long-read sequencing have made chromosome-scale genome assemblies possible for non-model organisms, (Feldmeyer et al., 2024). This includes many insects, which constitutes up to 75-80% of known animal species and of which roughly 80% remain undescribed (Gebremariam, 2024); yet high-quality annotation references remain concentrated in a handful of insect model species. Annotation is what makes a genome assembly applicable, and functional annotation is the harder half, since assigning a function relies predominantly from previously characterised genes. Predicted protein structures offer additional resources, as structural databases now cover much of known single-protein and complexes (Varadi et al., 2024; Abramson et al., 2024), and as a protein’s function is often defined by the partner it binds, a predicted complex can resolve gene families where sequence alone is insufficient to annotate. This is essential in non-model insects, whose annotation relies on gene orthologs from distant references such as *Drosophila melanogaster*.

In insects, neuropeptides are the most abundant and structurally diverse class of signalling molecules (Nässel and Zandawala, 2019). They are short secreted peptides roughly less than 40 amino acids long, produced and released by neurons and neuroendocrine cells to act as neurotransmitters, neuromodulators, and circulating hormones. Each is cleaved from a larger precursor and frequently carries a post-translational modification, such as C-terminal amidation, N-terminal pyroglutamate formation, or a disulfide bond, before becoming bioactive (Hook et al., 2018). Almost all act by binding a receptor at the target-cell surface, and most such receptors are G protein-coupled receptors (GPCRs), dominated by the rhodopsin-like (class A) and the secretin-like (class B) families (Caers et al., 2012).

The two spotted-cricket *G. bimaculatus* is an established laboratory model for hemimetabolous insects, studied in development, regeneration, neuroscience, and behaviour, and is increasingly farmed as food and feed (Horch et al., 2017). Feeding and metabolism, among the processes most strongly modulated by neuropeptides (Nässel and Zandawala, 2019), are especially well characterised in cricket, including the role of adipokinetic hormone signalling in dietary preference (Fukumura and Nagata, 2017; Fukumura et al., 2018). Recent genome assemblies now provide high sequencing quality, which make it possible to enrich a complete neuropeptide and receptor catalogue (Li et al., 2026; Kataoka et al., 2026; Ylla et al., 2021). Yet its neuropeptide and receptor complements remain only partially annotated, where the most complete catalogues comprise 41 neuropeptide families, and several sequences are still partial (Mochizuki et al., 2023; Kataoka et al., 2026). Closely related GPCR paralogues can have highly similar overall sequences but differ in ligand specificity. Receptors for species-specific peptides are difficult to assign by sequence similarity alone, since it requires a characterised homolog that non-model insects often lack (Shiraishi and Satake, 2026). Exhaustive deorphanization assays are therefore impractical, and ranking candidate pairings computationally may accelerate the screening.

Structure prediction now extends from single chains to complexes, so a neuropeptide and a candidate receptor can be folded together and their interface examined by residue. Folding each pair with two independently developed models, AlphaFold3 and Boltz-2 (Abramson et al., 2024; Passaro et al., 2025), allows the predictions to be cross-checked and evaluated with interface-restricted scores, ipSAE (Dunbrack, 2025) and interface-pLDDT, which summarize the reliability of the predicted contact rather than the receptor fold as a whole. We then cross-validate the receptor candidates for each family in a receptor phylogeny built from sequences of known ligand specificity. Because these models are trained on multiple sequence alignments, their confidence in an interface may partly reflect the evolutionary signals shared among related receptors. Applying this approach requires correct gene models, as fragmented or mispredicted sequences may fold the wrong protein. We therefore curated the neuropeptide and receptor sets by recovering truncated or missing models from automated gene finding, and validating the transmembrane topology across the receptor set. The resulting interactome, annotation, and curated sets are openly available through CricketBase (cricket.annotation.jp), which pairs a genome browser with a three-dimensional viewer of the predicted complexes, and the workflow offers a template for other non-model insects.

## Materials and Methods

A pipeline combining sequence- and structure-based evidence was designed to predict neuropeptide–receptor pairings in the two-spotted cricket (*Gryllus bimaculatus*) and to make the results available through a public genome resource, CricketBase (cricket.annotation.jp). Starting from the chromosome-level genome assembly, the re-annotation with BRAKER4 supplies protein models to two parallel gene discovery: one recovering and validating the G-protein-coupled receptor, and the other compiling and curating the mature neuropeptide set. The two parts merge at a combinatorial structure-prediction and interface-scoring stage that ranks every receptor–peptide pair. Phylogenetic analysis then confirms the top-ranked receptor candidates. The workflow is summarised in Figure 1 and detailed in the following sections.

**Figure 1.**
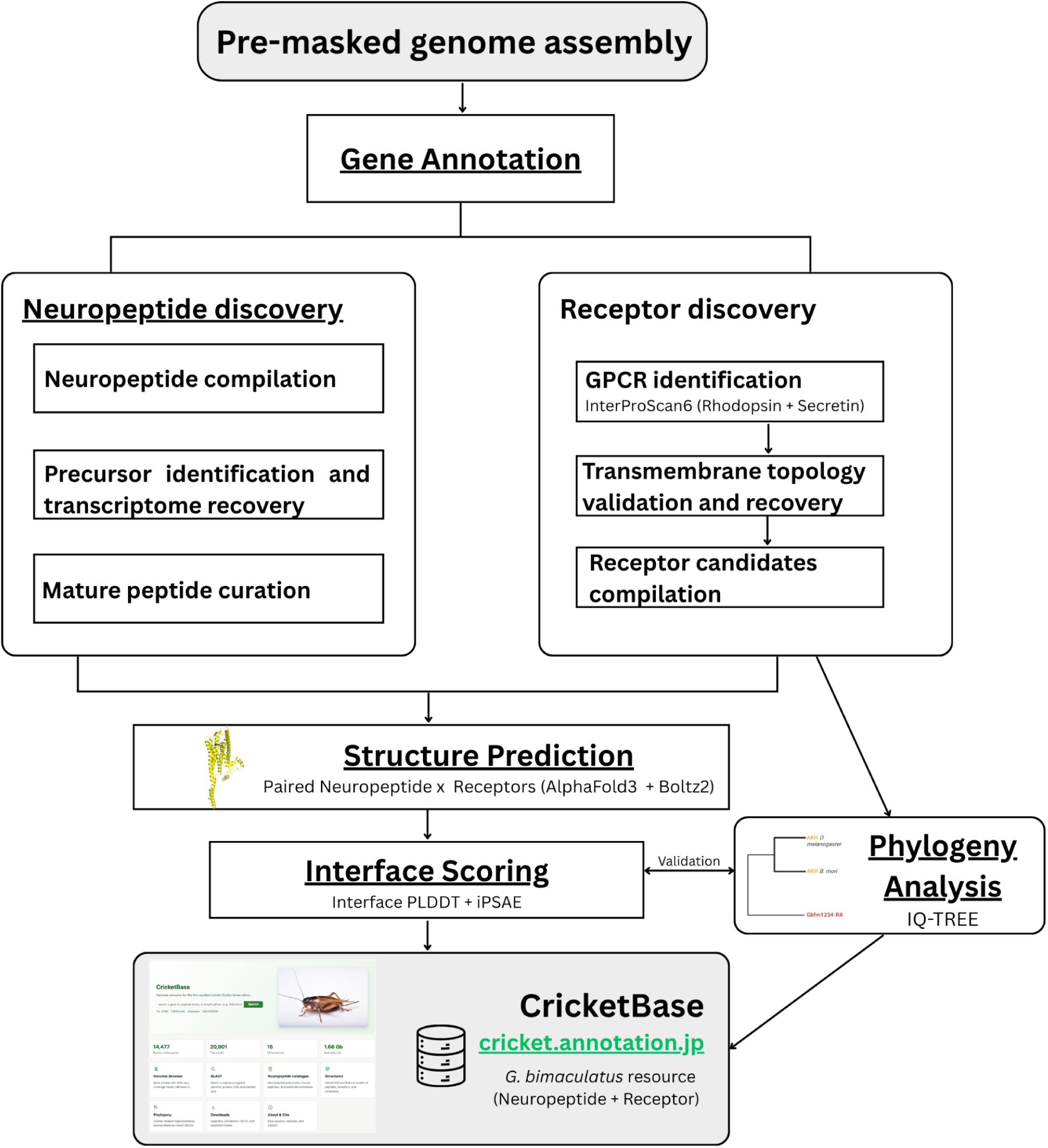
Sequence- and structure-based pipeline for annotating neuropeptide–receptor pairings in *Gryllus bimaculatus*.

**Figure 2.**
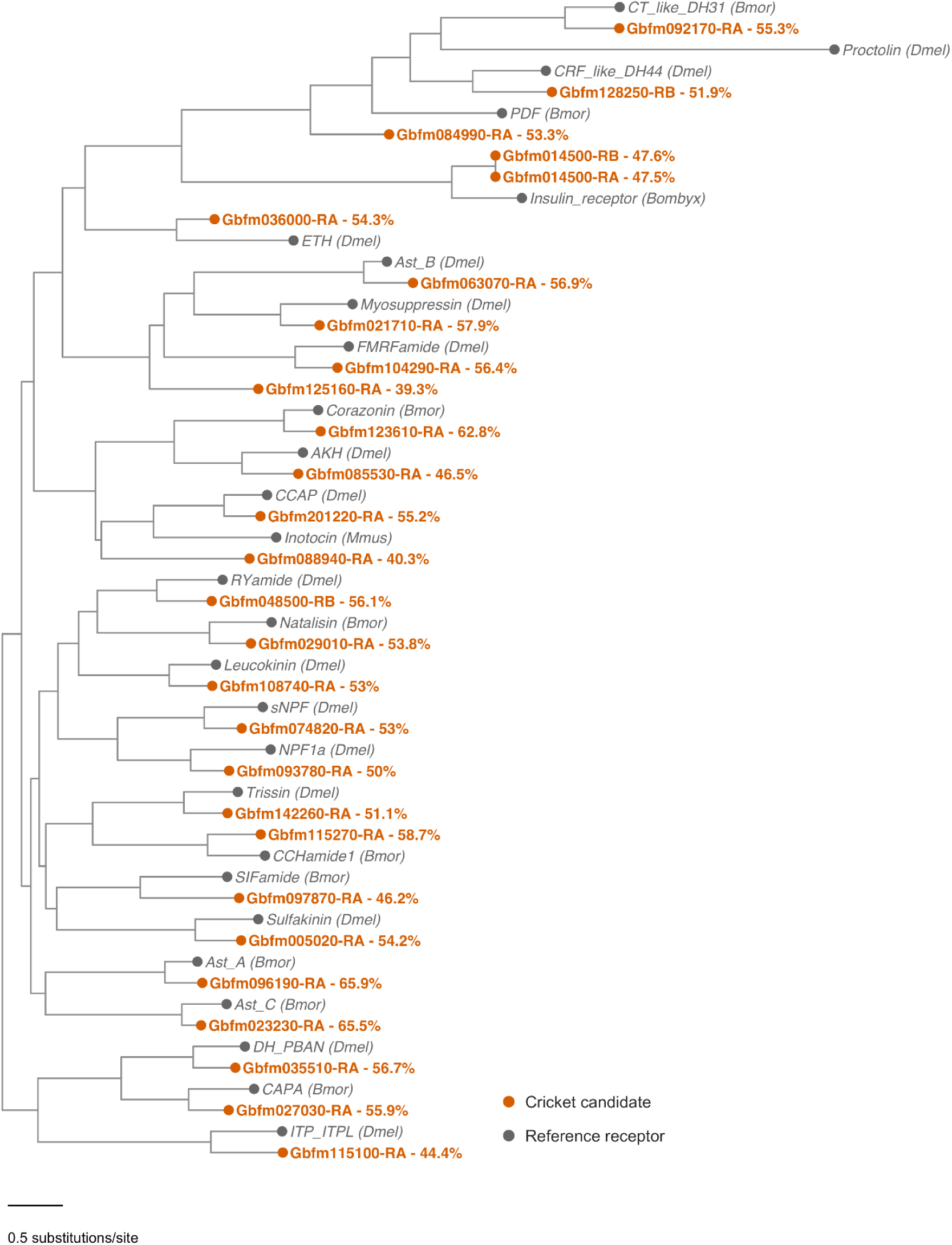
Receptor phylogeny of cricket neuropeptide-receptor candidates. Maximum-likelihood tree of the close phylogeny *G. bimaculatus* candidate receptors with reference receptors of known ligand specificity.

A chromosome-level genome assembly (Li et al., 2026) was re-annotated with BRAKER4 using protein and RNA-seq evidence. The resulting proteome corresponds to two discovery parts, neuropeptide (left) and receptor (right). The neuropeptide part recovers precursors from a curated reference set (Mochizuki et al., 2023) and defines mature peptides and their post-translational modifications. The receptor part identifies rhodopsin-family (A) and secretin-family (B) G-protein-coupled receptor candidates from protein-domain signatures, validates their seven-transmembrane topology, and recovers truncated or missing models from the transcriptome and from a second assembly (Kataoka et al., 2026) to give a non-redundant receptor set. Both of them merged together where every receptor is paired with every mature peptide, modelled with AlphaFold3 and Boltz-2, and ranked by interface score. The listed receptor candidates are then placed in a maximum-likelihood receptor phylogeny.

### Genome assemblies retrieval and annotation

Two *G. bimaculatus* chromosome-level assemblies were annotated, the primary assembly (Li et al., 2026) and a second assembly (Kataoka et al., 2026). The primary assembly was already soft-masked and was used directly; the second was repeat-masked with RepeatModeler2 and RepeatMasker before annotation. Both of the assemblies were annotated using BRAKER4 (Gabriel et al., 2024) with both RNA-seq and protein evidence from the Arthropoda partition of OrthoDB v12 (Tegenfeldt et al., 2025). The primary annotation was used without re-masking, where the secondary assembly was repeat-masked with RepeatModeler2 and RepeatMasker before annotation. RNA-seq evidence was sixteen paired-end Illumina libraries, seven spanning developmental stages (SRR14026720–SRR14026726) and nine from dissected body parts (DRR358356–DRR358364). Annotation completeness was assessed with BUSCO v6.1.0 against the OrthoDB v12 Insecta and Arthropoda lineages. Predicted proteomes were functionally annotated for their protein domains and family signatures with InterProScan 6 under its Nextflow workflow (Blum et al., 2026).

### Neuropeptide precursor identification and recovery

A reference set of neuropeptide precursors was compiled from the previous *G. bimaculatus* neuropeptide catalogue (Mochizuki et al. 2023) and extended with additional insect neuropeptide families from DINeR (Yeoh et al., 2017). Mature peptides from this set were searched with BLASTP against the BRAKER4 proteome to identify precursors already predicted by BRAKER4. Precursors not fully recovered by BLASTP were reconstructed with a transcriptome-guided procedure on the primary assembly. RNA-seq reads from the sixteen libraries were aligned to that assembly with HISAT2 (Kim et al., 2019), and transcripts were assembled with StringTie (Kovaka et al., 2019). Open reading frames were predicted with TransDecoder (Haas et al., 2013), with homology rescue against the precursor set and a reduced minimum ORF length of 50 aa appropriate for short neuropeptide precursors, and miniprot (Li, 2023) guiding the protein to genome alignment. Precursors recovered by BLASTP against the BRAKER4 proteome and precursors recovered by the transcriptome-guided approaches were combined into the final precursor set, each at its own genomic locus.

### Mature peptide curation and post-translational modifications

Putative mature peptide boundaries were inferred from the precursor sequences using known prohormone-processing patterns. Dibasic cleavage motifs, including KR and RR, and context-supported monobasic Arg sites were used as guides to identify candidate mature peptide regions (Veenstra, 2000). For mature peptides predicted to undergo C-terminal amidation, the terminal glycine in the precursor was treated as the amidation donor rather than as part of the final mature peptide sequence. SignalP 6.0 (Teufel et al., 2022) was used to predict the N-terminal secretory signal peptide and its cleavage site. Mature peptide boundaries and family assignments were manually curated and reviewed by insect neuropeptide experts. Ambiguous cases were resolved by comparison with conserved amino acid motifs characteristic of known mature peptides in the same neuropeptide family. When repeated peptide motifs separated by putative cleavage sites were present within a precursor, the arrangement of these repeats was also used to infer the mature peptide boundaries. Post-translational modifications relevant to the mature peptide properties, and to downstream structure prediction, were annotated for each peptide, covering pyroglutamate, amidation, cysteine disulfide bonds, and N-linked glycosylation.

### Neuropeptide receptor candidates discovery

Neuropeptide receptor candidates were extracted by matching curated signature sets for the two relevant GPCR classes retrieved from the predicted functional gene annotation by InterProScan6. Rhodopsin-like (class A) candidates were defined by Pfam PF00001 (7 transmembrane receptor (rhodopsin family)) or InterPro IPR017452 (GPCR, rhodopsin-like, 7TM) and IPR000276 (G protein-coupled receptor, rhodopsin-like). Secretin-like (class B) candidates were defined by PF00002 (7 transmembrane receptor (Secretin family)) or InterPro IPR000832 (GPCR, family 2, secretin-like) and IPR017981 (GPCR, family 2-like, 7TM). Proteins carrying these functional ID gene annotations were retained as candidate receptors of that class.

Candidate receptors were then screened for canonical seven-transmembrane (7TM) topology with DeepTMHMM (Hallgren et al., 2022), of which candidates predicted with seven transmembrane helices passed into the list. For candidates that did not show a complete 7TM topology, gene models were reconstructed using an approach independent of BRAKER4 to determine whether complete 7TM receptor models could be recovered. First, transcript models were assembled with StringTie using the RNA-seq alignments from all sixteen libraries, seven developmental-stage libraries (SRR14026720–SRR14026726) and nine dissected-body-part libraries (DRR358356–DRR358364). Open reading frames (ORFs) within these transcript models were then predicted with TransDecoder. In parallel, the protein sequences of receptor candidates lacking a complete 7TM topology were aligned to the genome with miniprot. This homology-based evidence was compared with the StringTie and TransDecoder models to determine whether a complete receptor model with the expected 7TM topology could be reconstructed at the corresponding locus. Using this procedure, full-length receptor models were recovered for genes that had been truncated or split into multiple fragments in the initial BRAKER4 annotation. A small number of candidates carried an over-extended N-terminal extension that gave a longer-than-canonical transmembrane topology. These models were corrected to the 7TM form by manual curation, trimming to the first valid methionine within a region of strong RNA-seq coverage.

To identify receptors present in the secondary assembly (Kataoka et al., 2026) but absent from the primary candidate set, the gene models were first generated using the same BRAKER4 evidence mode. Following that, the gene models with the same class-specific receptor candidates from InterProScan6 output were retrieved as candidate receptors. The candidate sets from the two assemblies were then clustered together with CD-HIT at 95% identity (−c 0.95) and a 50% coverage floor on the shorter sequence (−aS 0.5) (Fu et al., 2012) to merge partial or fragmented gene models in cross-assembly.

Each such additional receptor was mapped into the primary assembly coordinate system by combining homology-based localisation with transcript- and homology-guided gene modelling. First, the additional receptor’s protein sequence was searched against the primary assembly with TBLASTN to identify the candidate genomic locus. Following that, the same transcriptome-guided recovery from primary assembly was also applied to the secondary assembly. The compiled, topology-validated receptor set for each class, combining primary-assembly receptors, primary-mapped receptors recovered from the secondary assembly, and external secondary-assembly sequences, was compiled to non-redundant representatives for structure prediction by clustering with CD-HIT at the same parameters (−c 0.95 −aS 0.5) and taking each centroid as its representative, resulting in 66 rhodopsin- and 68 secretin-classes representatives.

### Structure prediction of peptide–receptor complexes

Each of the rhodopsin- and secretin-class receptor representatives with seven-transmembrane topology was paired with all 119 mature peptides *(Supplementary Table S2)*, giving 15,946 receptor-peptide combinations, and each pair was modelled independently with both AlphaFold3 v3.0.1 (Abramson et al., 2024) and Boltz-2 (Passaro et al., 2025). For consistency across all predictions and downstream analyses, in every input, the receptor was assigned to chain A and the mature peptide to chain B; for heterodimer peptide ligands (e.g., GPA2/GPB5, Bursicon-alpha/beta), where GPB5 and Bursicon-beta as the second peptide chain were assigned to chain C. Multiple sequence alignments (MSAs) were generated once per receptor and per peptide by the AlphaFold3 MSA workflow, and its json derived output was converted into a3m format with af3tools (https://github.com/cddlab/alphafold3_tools) to supply MSA information for Boltz-2. Through curation with insect neuropeptide experts, post-translational modifications were defined for the structure prediction to better represent the binding interaction of the mature peptide. The modifications were presented for the structural inference of the structure prediction models, as the MSA searches only the unmodified mature peptide sequence, where each mature peptide could carry none, one, or several of the modification types. N-terminal pyroglutamate, formed by cyclisation of an N-terminal glutamine or glutamate, was represented as the residue modification PCA. C-terminal amidation is assigned to peptides with C-terminal glycine and dibasic (or relevant monobasic) cleavage motif and was added as an NH2 ligand bonded to the terminal carbonyl carbon through an explicit bonded-atom pair. An intramolecular disulfide bond was specified as an SG-SG bond between the relevant cysteine pair. N-linked glycosylation was modelled as N-acetylglucosamine stub bonded from the asparagine side chain to the glycan at the mapped sequon.

### Interface confidence scoring for candidate receptor shortlisting

Every predicted complex was scored at the receptor-peptide interface with two metrics that measure different structure properties. Interface pLDDT is a local measure of the model’s per-residue confidence at the contact surface, on a 0 to 100 scale. AlphaFold3 reports pLDDT per atom, and these values were averaged within each residue; Boltz-2 reports one value per residue, which was matched to the residue by chain and residue number. On the other hand, ipSAE is a relational measure of how confidently the two chains are positioned relative to each other, on a 0 to 1 scale, derived from the predicted aligned error (PAE), the model’s estimate of positional error between residue pairs. Both scoring were calculated from the AlphaFold3 and Boltz-2 outputs, pLDDT from the model mmCIF structure file and PAE from their corresponding confidence files.

Interface pLDDT is the mean per-residue confidence over the residues that form the receptor-peptide contact. AlphaFold3 reports pLDDT per atom, so these values were averaged within each residue; Boltz-2 reports one value per polymer residue directly. A residue was counted as part of the interface when any of its atoms lay within 10 Å of an atom in the partner chain. Two summaries were computed from the per-residue values: a size-weighted mean over all interface residues pooled across both chains (*Supplementary Methods Equation S1d*), and a balanced mean that averages the two chain-level interface means (*Supplementary Methods Equation S1e*). The balanced form was used for ranking as the receptor itself is longer relative to a short peptide, so the size-weighted mean is dominated by the many receptor interface residues scoring, whereas the balanced form gives peptide and receptor equal weight.

ipSAE is the mean, over residue pairs spanning the two chains, of a TM-score-like (template-modelling score) function of their predicted aligned error. A pair contributed to the score when its inter-chain PAE fell below 10 Å (*Supplementary Methods Equation S2b*), so the residues entering ipSAE are selected by predicted positional error rather than by physical distance. The normalisation length *d*_0_ is set from the number of partner residues passing the cutoff rather than from full chain length (*Supplementary Methods Equation S2c*), so the score reflects the actual formed interface rather than the size of the complex protein structure. Because predicted aligned error is not symmetric between chains, this calculation gives a different value depending on which chain is treated as the reference: one value scores chain A against chain B, the other scores chain B against chain A. The higher score between these two is the one used for listing the putative receptor candidates.

For each mature peptide, complexes were ranked separately by ipSAE and by balanced interface pLDDT, and three shortlists were produced: the top five rhodopsin-class receptors, the top five secretin-class receptors, and the top ten receptors across both classes.

### Phylogeny mapping of known neuropeptide receptors

A reference set of 76 neuropeptide receptors from different species across our known 32 neuropeptide families was collected from NCBI RefSeq and UniProt (Bateman et al., 2022), guided by prior receptor curation (Hewes and Taghert, 2001). All of the neuropeptide receptor candidates were placed in a maximum-likelihood phylogeny with reference receptors of known ligand specificity. Most curated known receptor sequences come from *Drosophila melanogaster* and *Bombyx mori*, with other sequence entries from *Bombus terrestris*, *Aedes aegypti*, *Homo sapiens*, and *Mus musculus*. Closely related families and paralogues were kept in the same alignment for the receptor curation assignment (e.g., sNPF with NPF1a and NPF1b, CCHamide-1 with CCHamide-2, and Ast C with Ast CC and Ast CCC). Sequences were then aligned with MAFFT v7.526 (Katoh and Standley, 2013) and trimmed with trimAl v1.5.rev1 (Capella-Gutierrez et al., 2009) to remove gap-rich and ambiguously aligned positions, as receptor alignments carry long N-terminal and loop segments that align poorly across families. Columns with residues in fewer than 20 percent of sequences were discarded (-gt 0.2), and at least 40 percent of the original columns were retained (-cons 40). Maximum-likelihood trees were generated with IQ-TREE 3.1.2 (Wong et al., 2026) using 1000 ultrafast bootstrap replicates (Hoang et al., 2018) and 1000 SH-aLRT replicates (Guindon et al., 2010). The candidate tree was inferred under the LG+F+G4 substitution model. A receptor was retained as a phylogenetically supported candidate member of a family when it grouped with that family’s reference receptor at SH-aLRT >= 80 and ultrafast bootstrap >= 95. Ultrafast bootstrap measures how often a clade is recovered from resampled alignments and SH-aLRT tests the branch against its collapse to zero length.

## Results

### Re-annotation of the reference assembly

BRAKER4, run in ortholog-protein and RNA-seq evidence mode, produced 14,477 protein-coding genes and 20,901 transcripts in the genome assembly. Proteome BUSCO completeness was 95.3% against the Insecta OrthoDB v12 set (n = 3,114) and 99.3% against Arthropoda OrthoDB v12 (n = 1,667), both higher than the published annotation of the same assembly, at 86.7% and 92.4% respectively (Table 1). To assess how many previously reported gene models this re-annotation recovered, we compared the new gene set against the published annotation of the same assembly and found that 11,152 of the 14,457 published gene models were recovered (77.1%).

**Table 1.** BRAKER4 re-annotation compared with the published annotation from prior assembly annotation (Li et al., 2026), with BUSCO completeness scored against OrthoDB v12 lineage datasets of Insecta and Arthropoda.

| Metric | Previous annotation | This study |
| --- | --- | --- |
| Protein-coding genes | 14,457 | 14,477 |
| Transcript (mRNAs) | 19,058 | 20,901 |
| Proteome BUSCO<br>(insecta_odb12) | 86.7% | 95.3% |
| Proteome BUSCO<br>(arthropoda_odb12) | 92.4% | 99.3% |

**Table 2.** The 48 curated neuropeptide families of *G. bimaculatus* (51 precursor loci).

| No. | Neuropeptide | Gene ID |
| --- | --- | --- |
| 1 | ACP | Gbfm010640-RA |
| 2 | AKH | Gbfm010690-RA |
| 3 | Agatoxin-like peptide | Gbfm064660-RA |
| 4 | Allatotropin | Gbfm054560-RA |
| 5 | Ast A | Gbfm018390-RA, Gbfm018390-RB |
| 6 | Ast_B | Gbfm200000-RA |
| 7 | Ast_C | Gbfm200010-RA |
| 8 | Ast_CC | Gbfm087330-RA |
| 9 | Ast_CCC | Gbfm126330-RA |
| 10 | Ast_CCC | Gbfm126330-RB |
| 11 | Bursicon_alpha | Gbfm101180-RA |
| 12 | Bursicon_beta | Gbfm101190-RA |
| 13 | CAPA | Gbfm018360-RA |
| 14 | CCAP | Gbfm051860-RA |
| 15 | CCHamide1 | Gbfm107500-RA |
| 16 | CCHamide2 | Gbfm107490-RA |
| 17 | CNMamide | Gbfm092720-RA |
| 18 | CRF-like DH44 | Gbfm105870-RA |
| 19 | CT-like DH31 | Gbfm018140-RA |
| 20 | Corazonin | Gbfm120730-RB |
| 21 | DH/PBAN | Gbfm030670-RA, Gbfm200020-RA, Gbfm018360-RA |
| 22 | ETH | Gbfm002910-RA |
| 23 | Eclosion_hormone | Gbfm122320-RA |
| 24 | Elevenin | Gbfm041140-RA |
| 25 | FMRFamide | Gbfm020420-RC |
| 26 | GPA2 | Gbfm004320-RA |
| 27 | GPB5 | Gbfm000500-RA |
| 28 | IDLSR-like peptide | Gbfm028170-RA |
| 29 | ITG-like peptide | Gbfm200040-RA |
| 30 | ITP/ITPL | Gbfm142070-RC |
| 31 | Inotocin | Gbfm140540-RA |
| 32 | Insulin-like peptide | Gbfm200030-RA |
| 33 | Leucokinin | Gbfm004480-RA |
| 34 | Myosuppressin | Gbfm025500-RA |
| 35 | NPF1a | Gbfm200050-RA |
| 36 | NPF1b | Gbfm106950-RA |
| 37 | NPLP1 | Gbfm125890-RA |
| 38 | NPP1 | Gbfm111050-RA |
| 39 | NPP2 | Gbfm050980-RA |
| 40 | Natalisin | Gbfm004490-RA |
| 41 | Orcokinin | Gbfm077450-RA |
| 42 | PDF | Gbfm086660-RA |
| 43 | Proctolin | Gbfm009610-RA |
| 44 | RYamide | Gbfm003930-RA |
| 45 | SIFamide | Gbfm002920-RA |
| 46 | Sulfakinin | Gbfm083320-RA |
| 47 | sNPF | Gbfm108190-RA |
| 48 | Trissin | Gbfm032990-RA |

### Curated neuropeptide precursor families

A total of 48 neuropeptide families, referencing to the compiled dataset from the reference precursors from existing *G. bimaculatus* neuropeptide catalogue (Mochizuki et al., 2023) and DINeR (Yeoh et al., 2017), were discovered through gene models set from BRAKER4 and transcriptome recovery. 43 neuropeptide families were successfully identified through the gene models set, whereas 5 neuropeptide families were recovered through reconstruction from RNA-seq evidence. From a total of 48 neuropeptide families, seven precursors are newly identified or completed for *G. bimaculatus*: agatoxin-like peptide, ITG-like peptide, NPLP1, CNMamide, IDLSR-like peptide, inotocin, and natalisin. Natalisin had previously been reported only as a partial cDNA hit, and it is now resolved to its full sequence and complete mature peptide. Curation of mature peptides through insect experts, known neuropeptide patterns, and SignalP confirmation of secretory signals and cleavage sites yielded 119 distinct mature peptides, with post-translational modifications (pyroglutamate, C-terminal amidation, disulfide bonds, and N-glycosylation) annotated for structure modelling.

### Curated neuropeptide receptor candidates

Functional-based extraction from the final InterProScan 6 annotation gave 113 rhodopsin-class and 92 secretin-class GPCR candidates in the primary assembly (Supplementary Table S2). Of these, 60 rhodopsin-class and 61 secretin-class carried a complete seven-transmembrane topology as annotated, and a further 13 and 19 returned a complete topology after reconstruction. Three candidates, two rhodopsin-class and one secretin-class, were trimmed on its first methionine from the over-extended with an eighth helix from an extended N terminus. The remaining 38 rhodopsin-class and 11 secretin-class candidates reached no complete topology and were not processed for structure prediction. The secondary assembly further added the candidates, where 9 rhodopsin and 11 secretin sequences with seven helix were added to the candidate list. Six of each from secondary assembly, rhodopsin and secretin, were able to be placed in the primary assembly, where the remaining three rhodopsin and five secretin sequences were marked as external gene models. The compiled set comprised 81 rhodopsin-class and 92 secretin-class 7TM receptors, resulting in a total of 66 rhodopsin-class and 68 secretin-class representatives, used for structure prediction.

### A structure-based candidate interaction map with phylogeny validation

Ranking the 15,946 scored complexes by ipSAE and by balanced interface pLDDT produced a top-five rhodopsin, top-five secretin, and top-ten combined shortlist for each of the 119 mature peptides. Placing the top-scoring candidate for each family in the receptor phylogeny, most of the candidates fell within a clade that also contained at least one reference receptor. For 21 of the 27 distinct receptors, the candidate grouped with its own family’s reference receptor at strong support, most clearly for the allatostatin-A, allatostatin-C, and corazonin receptors, where the candidate occupied a short branch adjacent to its *Drosophila* or *Bombyx* ortholog. Of the remaining six, five grouped with their own family’s reference at lower support: AKH, ETH, and the insulin-like receptor at a weakly supported node, and inotocin and PDF only within a larger clade that also contained other families. Proctolin was the exception, its candidate not grouping with the proctolin reference at any level of the tree. The 66 rhodopsin and 68 secretin representatives were paired with all 119 mature peptides, and the 15,946 resulting complexes were each modelled with both AlphaFold3 and Boltz-2 and scored at the interface. For every mature peptide, ranking by ipSAE and by balanced interface pLDDT produced a top-five rhodopsin shortlist, a top-five secretin shortlist, and a top-ten combined shortlist of the most confidently predicted receptor partners. The results of the phylogenetic analysis strongly supported those of the structural analysis, and those without known neuropeptide receptors can be narrowed down with structural screening.

### A reproducible resource for neuropeptide - receptor research

The genome annotation, recovered gene models, curated neuropeptide and receptor sets, mature-peptide set, and ranked predicted complexes are openly available through CricketBase (https://cricket.annotation.jp), which combines a genome browser from JBrowse2 (Diesh et al., 2023) with a protein structure visualization from NGL (Rose et al., 2018) (Figure 3). The genome browser (Figure 3A) shows the gene models, the recovered receptor and neuropeptide loci, and BigWig coverage tracks from the RNA-seq alignments, and a genome map (Figure 3C) locates every neuropeptide precursor and 7TM receptor across the 15 pseudochromosomes. The structure explorer (Figure 3D) displays each predicted peptide-receptor complex alongside the standalone peptide and receptor models. It reports the top-model ipSAE and balanced interface pLDDT (Figure 3E), highlights the interface residues on the structure, and links each receptor candidate to its transmembrane topology from DeepTMHMM, its InterProScan domains, its SignalP-confirmed signal peptide (Figure 3B), and its genomic locus. A phylogenetic tree (Figure 3F) places the candidate receptors among known neuropeptide receptors from other species. Altogether, the integrated genome, structure, and phylogeny information connect each neuropeptide to its ranked candidate receptors and the associated confidence scores, phylogenetic placement, and domain annotation, providing a basis for prioritising deorphanization experiments.

**Figure 3.**
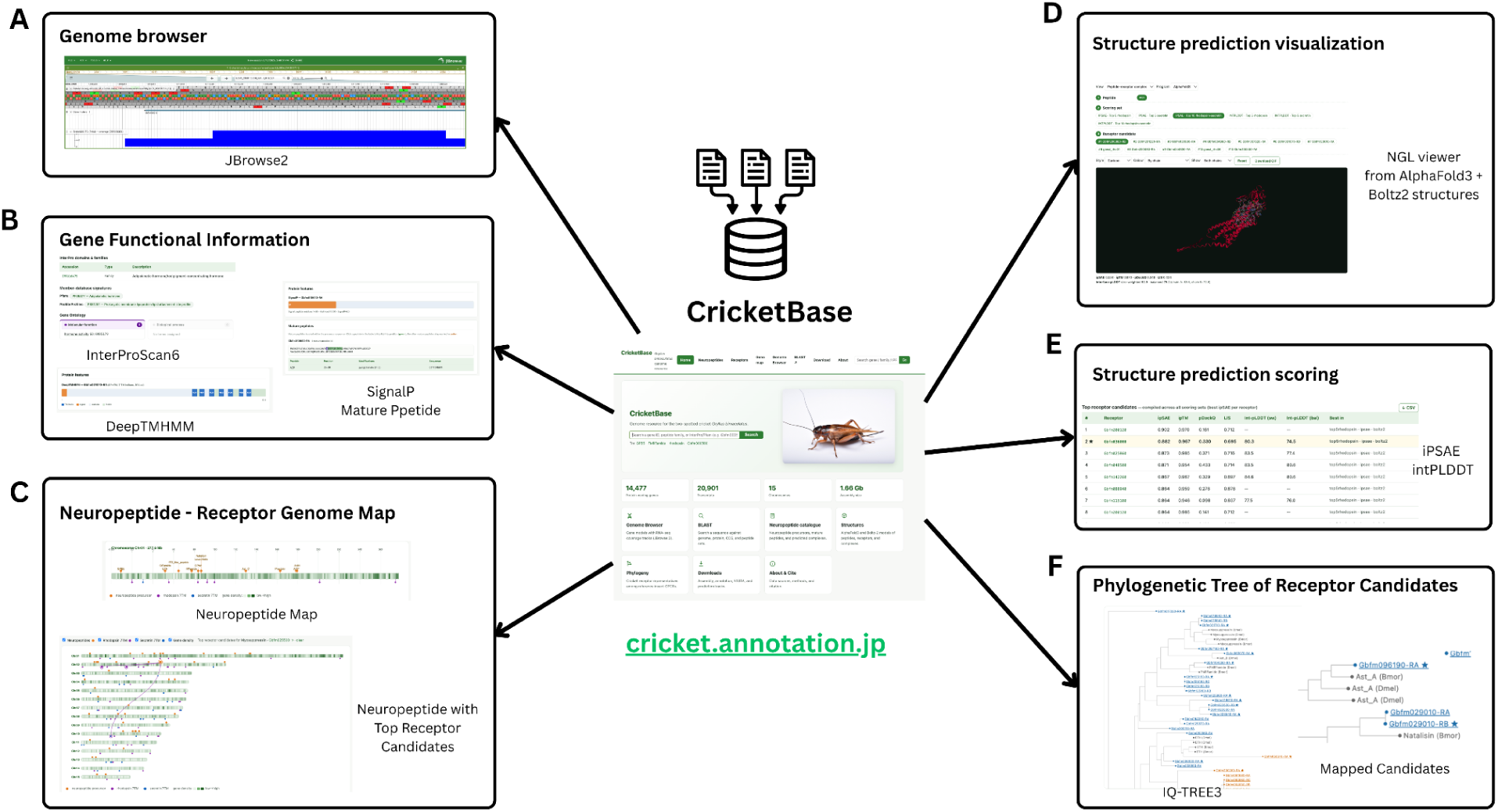
CricketBase, an integrated genome, structure, and phylogeny resource for G. bimaculatus neuropeptide-receptor research. (A) JBrowse2 genome browser with gene models, recovered neuropeptide and receptor loci, and RNA-seq coverage tracks. (B) Gene functional information, including InterProScan domains and families, DeepTMHMM transmembrane topology, and SignalP signal-peptide prediction with the curated mature peptides. (C) Genome map of neuropeptide precursor and 7TM receptor positions across the chromosomes, with the top-ranked receptor candidates for each neuropeptide shown against gene density. (D) NGL structure viewer for the AlphaFold3 and Boltz-2 peptide–receptor complexes. (E) Ranked interface-confidence scores, ipSAE and balanced interface pLDDT, for the receptor candidates of a single peptide. (F) Maximum-likelihood receptor phylogeny placing the cricket candidates among reference receptors of known ligand specificity, with mapped candidates marked.

## Discussion

### Curation recovers loci and corrects models that automated annotation misses

The annotation workflow recovered peptides and corrected receptor models that automated pipelines missed, truncated, or extended incorrectly. Of the 51 curated neuropeptide precursor loci, 24 of them carry no functional annotation from InterProScan6, and may not have been identified from domain annotation alone. The CNMamide precursor (Gbfm092720) is one such case, as it is recovered through our workflow by direct comparison with known CNMamide precursors and predicted signal peptide confirmation. Nevertheless, a functional domain hit may also refer to a wrong family, as such, the ACP (adipokinetic hormone/corazonin-related peptide) precursor (Gbfm010640) carries a Pfam hit assigned to the adipokinetic hormone/red pigment-concentrating hormone family. Without the neuropeptide curation workflow, ACP may have been misclassified as AKH (adipokinetic hormone).

In the same way of misclassified neuropeptide from automated gene annotation, the candidate receptor set also required correction. Of 119 rhodopsin-class and 98 secretin-class receptor candidates, only 78 and 87, respectively, carried a complete seven-transmembrane topology as originally annotated. Therefore, we provided reannotation with additional gene-finding evidence to extend or trim the incorrect models to restore the receptor transmembrane topology. These corrections address a problem broader than just for the two-spotted cricket genome. Many neuropeptide receptors remain orphan even in well-studied insects, and the curated, ranked candidate list in CricketBase, built from structure prediction rather than sequence similarity alone, offers a starting point for deorphanization screens in *Gryllus bimaculatus* and, by the same workflow, in other non-model insects.

### Phylogenetic placement supports the structural receptor assignments

Both phylogenetic and structural analyses support the further receptor screening, as a clade reflects the descent from a common ancestor, whereas an interface score reflects whether AlphaFold3 or Boltz-2 can model a specific docking geometry with confidence. For most families, the candidate receptor that scored highest in the structural screen is also the one that groups with its own family’s reference receptor, most clearly for the allatostatin-A, allatostatin-C, and corazonin receptors (65.9%, 65.5%, and 62.8% identity), where each complex attains one of the highest interface-confidence scores among candidates (Supplementary Figure S2). Where phylogenetic support is weaker, a high structural score alone can still designate a candidate, and such assignments are treated as provisional to phylogeny information. A weakly supported node indicates limited resolution among closely related sequences in the reference set, not necessarily an incorrect receptor. One such exception is proctolin, where its candidate does not group close to the branch of its reference receptor, so this assignment depends on the structural score alone.

### Limitations and future outlook

Gene model curation for neuropeptides and their receptors still involves rigorous manual steps due to the short mature peptides and the genetic distance between *Gryllus bimaculatus* and the availability of close reference species. Domain-based annotation depends on a conserved region long enough to model as a profile, whereas identifying a mature neuropeptide requires the precursor context and the varied cleavage sites flanking a motif of a few residues, where information that automated annotation does not capture well. On the other hand, receptors are easier to recognise, since their sequence motifs are longer and the seven-transmembrane architecture is possible to predict. However, assigning a ligand to a specific receptor is still difficult due to their high sequence similarity in the candidate receptor set and gene model completeness from truncated or over-extended gene model predictions.

This neuropeptide-receptor interactome was therefore built primarily through manual curation grounded in transcriptome evidence and structure-prediction-based deorphanization screening, rather than by transferring gene models from a distant, well-annotated relative such as *Drosophila melanogaster* or *Bombyx mori*, recovering gene models that ortholog-based annotation alone would have missed. The same framework should extend to non-model insect counterparts that lack a close annotated reference, though scaling it to a broader range of genomes would require both transcriptome evidence for every target species and a classifier that learns neuropeptide precursor architecture and neuropeptide receptor transmembrane topology as a whole.

Structure prediction scoring itself complements the sequence evidence from phylogenetic mapping, and it’s especially useful for narrowing down candidates for peptides whose receptors are still orphan due to unknown reference. While pairing of all possible neuropeptide and receptor candidates is feasible for a single species, it becomes computationally extensive at a larger taxonomic scale, where the more practical direction is to prioritise candidate pairs before prediction rather than to predict every possible neuropeptide-receptor combination. In the future, we plan to incorporate this information to construct a machine learning model that can automate the discovery of neuropeptide precursors and their receptors, together with predicting novel neuropeptides. In addition, we aim to also provide a resource-optimized model with clear motif patterns that can efficiently screen neuropeptide and receptor pairing.

## Conclusion

We combined sequence-based annotation with structure-prediction scoring to characterise the neuropeptide–receptor interactome of *G. bimaculatus* as a comprehensive resource. Re-annotating a chromosome-scale assembly, curating the neuropeptide and rhodopsin- and secretin-class GPCR complements, and recovering fragmentary or missing models from the transcriptome and a second assembly produced a more complete catalogue than either genome offered alone, including seven newly identified or completed neuropeptides and the full neuropeptide of natalisin. Modelling every peptide against every receptor with two independent predictors and scoring each interface with pLDDT and ipSAE turned that catalogue into a ranked map of candidate pairings that can distinguish confidently docked complexes. A receptor phylogeny gave an independent check on the top-ranked candidates and agreed with the high structural scores for most families. These pairings are computational predictions and remain to be tested experimentally, but they provide a prioritised shortlist that directs deorphanization to its most plausible targets, and a curated foundation for dissecting neuropeptide control of feeding, metabolism, and reproduction in this model. Openly available through CricketBase (https://cricket.annotation.jp), the sequence- and structure-based approach of this study may extend to other difficult gene families, across non-model insects.

## Data Availability

Implemented code is available at https://github.com/frozuyuko/CricketBaseCode, and the output is available at https://cricket.annotation.jp

## Acknowledgments

We acknowledge the assistance of Claude in suggesting code for designing CricketBase web interface. The authors checked the database code, analysis code, and performed all manuscript editing.

